# Mu Opioid Receptor Gene Dosage Influences Reciprocal Social Behaviors and Nucleus Accumbens Microcircuitry

**DOI:** 10.1101/2020.08.24.265454

**Authors:** Carlee Toddes, Emilia M. Lefevre, Dieter D. Brandner, Lauryn Zugschwert, Patrick E. Rothwell

## Abstract

The mu opioid receptor regulates reward derived from both drug use and natural experiences, including social interaction. Homozygous genetic knockout of the mu opioid receptor (Oprm1−/−) causes social deficits in mice, whereas partial dysregulation of mu opioid signaling has been documented in several neuropsychiatric disorders. Here, we investigated the social behavior of male and female mice with heterozygous genetic knockout of the mu opioid receptor (Oprm1+/−), modeling partial reduction of mu opioid signaling. Reciprocal social interaction and social conditioned place preference were diminished in Oprm1+/− and Oprm1−/− mutants of both sexes. Interaction with Oprm1 mutants also altered the social behavior of genotypical test partners. We corroborated this latter result using a social preference task, in which genotypical mice preferred interactions with another typical mouse over Oprm1 mutants. We also analyzed inhibitory synapses in the nucleus accumbens, a key brain region for mu opioid regulation of social behavior, using methods that differentiate between medium spiny neurons (MSNs) expressing the D1 or D2 dopamine receptor. Inhibitory synaptic transmission was increased in D2-MSNs of male mutants, but not female mutants, while the density of inhibitory synaptic puncta at the cell body of D2-MSNs was increased in both male and female mutants. These changes in nucleus accumbens microcircuitry were more robust in Oprm1+/− mutants than Oprm1−/− mutants, demonstrating that partial reductions of mu opioid signaling can have large effects on brain function and behavior. Our results support a role for partial dysregulation of mu opioid signaling in social deficits associated with neuropsychiatric conditions.

## INTRODUCTION

Mu opioid receptor activation facilitates reward derived from social interaction and other natural experiences, as well as the abuse liability of exogenous opiate narcotics [1-3]. Agonists with high mu opioid receptor affinity increase visual attention to faces in humans, and enhance social play behavior in juvenile rodents as well as marmosets, while pharmacological blockade of opioid receptors causes deficits in these behaviors [4-6]. Mu opioid receptor availability in the human nucleus accumbens is regulated by a variety of social circumstances [7,8], and intra-accumbal manipulations of mu opioid receptor activation can bidirectionally modulate social behavior in rodents [9-11]. These findings are consistent with a general role for mu opioid receptor activation within the nucleus accumbens in motivated behavior [12-14].

Dysregulation of mu opioid receptor signaling may contribute to deficits in social interaction and other motivated behaviors that are a hallmark of neuropsychiatric disorders [15-19]. Mice with constitutive genetic knockout of the mu opioid receptor (Oprm1) have behavioral deficits in social affiliation, attachment, and reward, as well as dramatic remodeling of synaptic architecture and gene expression in the nucleus accumbens [20-22]. These studies have focused on homozygous knockout mice, but partial loss of mu opioid receptor function (as modeled by heterozygous Oprm1 knockout) is likely more relevant to functional deficits in human neuropsychiatric disorders. However, the influence of Oprm1 gene dosage on social behavior and nucleus accumbens function remains unclear.

To investigate these issues, we tested heterozygous and homozygous Oprm1 knockout mice on a battery of social behavior assays. To evaluate the quality of reciprocal social interaction, we also quantified the social behavior of wildtype partners paired with Oprm1 mutants during behavioral testing, and measured the preference of wildtype mice for social interaction with Oprm1 mutants. Our results suggests Oprm1 gene dosage influences social behavior of both mutant mice and interacting partners in a reciprocal fashion. These behavioral changes were accompanied by cell type-specific alterations in the structure and function of inhibitory synapses formed onto medium spiny neurons (MSNs) in the nucleus accumbens. This cell type- and synapse-specific remodeling of nucleus accumbens microcircuitry may contribute to the deficits in reciprocal social behavior caused by partial loss of mu opioid receptor signaling.

## MATERIALS AND METHODS

### Subjects

Experiments were performed with female and male C57BL/6J mice, Oprm1 knockout mice [23], Drd1a-tdTomato BAC transgenic mice [24], and Drd2-eGFP BAC transgenic mice [25]. Experimental procedures were approved by the Institutional Animal Care and Use Committee of the University of Minnesota. For additional details, see Supplementary Information.

### Gene Expression

Quantitative RT-PCR was performed as previously described [26]. Primer sequences for Oprm1 and beta-actin (control) can be found in Table S1. For additional details, see Supplementary Information.

### Behavioral Assays

Measurement of thermal antinociception and open field locomotion after morphine administration were performed as previously described [26]. To evaluate social behavior, we used a battery of previously described assays: social conditioned place preference [21,27,28]; reciprocal social interaction [29]; the standard three-chamber test of sociability and preference for social novelty [30]; and a real-time preference test for typical social interaction [31]. For additional details, see Supplementary Information.

### Electrophysiology

Whole-cell voltage-clamp recordings from nucleus accumbens MSNs in acute brain slices were performed as previously described [32]. Miniature inhibitory post-synaptic currents (mIPSCs) were recorded at a holding potential of 0 mV and pharmacologically isolated using D-APV (50 mM), NBQX (10 mM), and tetrodotoxin (0.5 µM). For additional details, see Supplementary Information.

### Immunohistochemistry

Tissue sections from Oprm1+/+, Oprm1+/−, and Oprm1−/− mice carrying Drd2-eGFP were incubated with antibodies against GFP (to label D2-MSN somata) and gephyrin (to label inhibitory synapses). Confocal image stacks from stained tissue sections were rendered in 3D and analyzed using Imaris 9.0 (Bitplane, Zurich, Switzerland). For additional details, see Supplementary Information.

### Statistical Analyses

Analysis of variance (ANOVA) was conducted in IBM SPSS Statistics v24, with details provided in Supplemental Information. In the text, we report significant effects that are critical for data interpretation, but comprehensive reporting of all main effects and interactions from ANOVA models can be found in Table S2. Significant main effects or interactions are denoted by black and red stars above the data, respectively. All summary data are displayed as mean ± SEM, with individual data points from female and male mice shown as open and closed symbols, respectively.

## RESULTS

### Functional Validation of Partial Genetic Knockout in Oprm1+/− Mutant Mice

To compare Oprm1+/− and Oprm1−/− mice with Oprm1+/+ littermates, we first studied female and male offspring generated by breeding two Oprm1+/− parents (Figure 1A). We used quantitative RT-PCR to validate complete loss of Oprm1 expression in brain tissue from Oprm1−/− mice, with a partial reduction of expression in Oprm1+/− mice (Figure 1B; F_2,11_=128.63, p<0.001), indicating a gene dosage effect. Following administration of morphine, Oprm1−/− mice did not exhibit dose-dependent increases in (1) hyperlocomotion in the open field (Figure 1C) or (2) thermal antinociception on the hot plate (Figure S1) [23]. In these assays, the behavioral response of Oprm1+/− mice were attenuated but not completely absent, consistent with previous publications [33] and the notion that both Oprm1 alleles contribute to expression of functional receptors [34].

**Figure 1.**
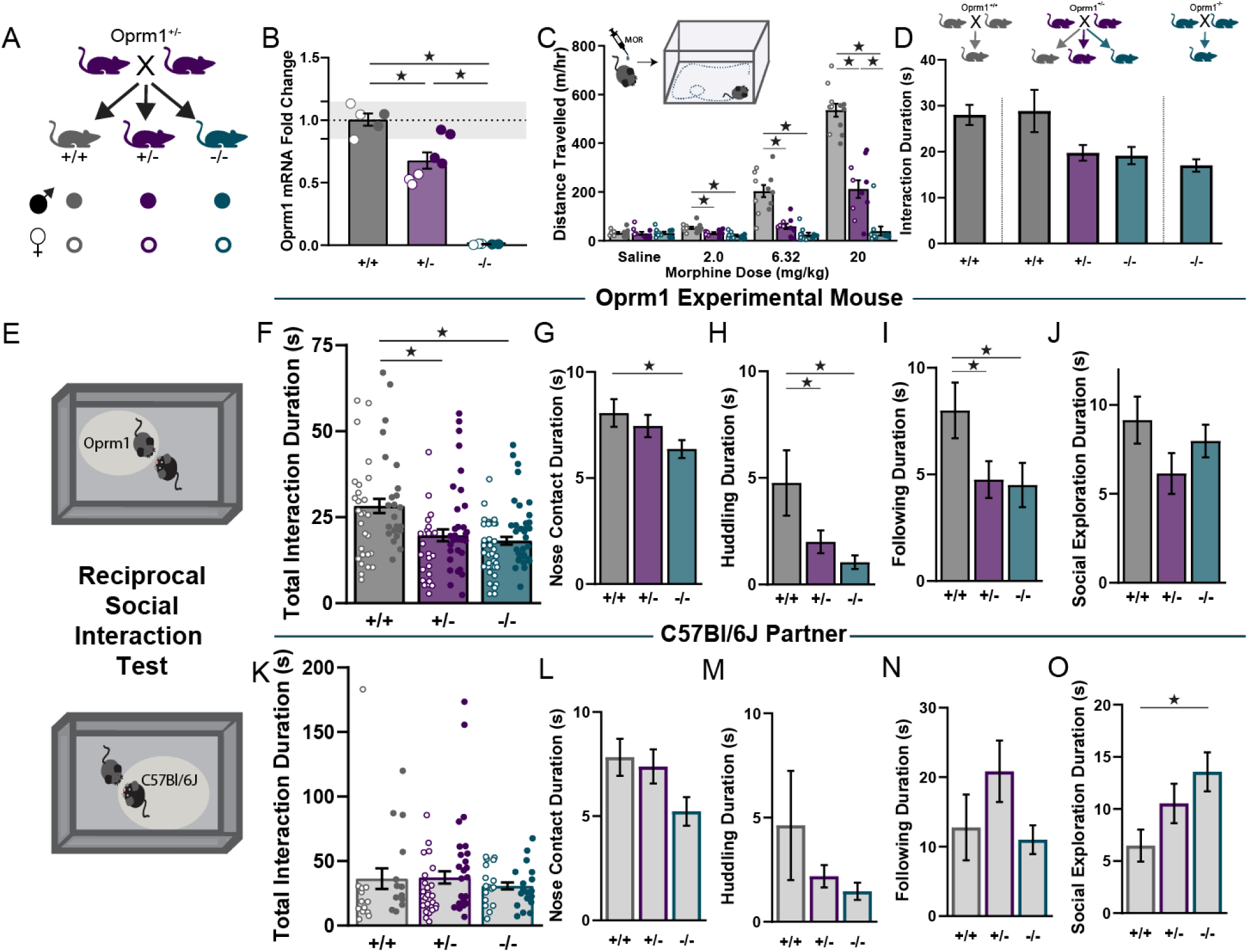
Validation of Oprm1 gene dosage and influence on reciprocal social interaction. **(A)** Breeding strategy used to generate littermates of all possible genotypes for validation experiments (top), with legend defining appearance of individual data points for each genotype and sex (bottom). **(B)** Assessment of mu opioid receptor (Oprm1) mRNA levels in striatal tissue punches using quantitative PCR. **(C)** Distance travelled in a test of open field activity after injection of morphine. **(D)** Breeding strategies used to generate mice for tests of social behavior (top), with reciprocal social interaction data disaggregated by breeding strategy (bottom). **(E)** Schematic diagram of the reciprocal social interaction test, separately highlighting behavior of the Orpm1 mutant mouse (top) and the genotypical testing partner (bottom). **(F-J)** Reciprocal social interaction durations for Oprm1 mutant mice: total interaction **(F)**, nose contact **(G)**, huddling **(H)**, following **(I)**, and social exploration **(J). (K-O)** Reciprocal social interaction durations for genotypical partners: total interaction **(K)**, nose contact **(L)**, huddling **(M)**, following **(N)**, and social exploration **(O)**. All groups contained similar numbers of female mice (open symbols) and male mice (closed symbols); see Supplementary Table 2 for detailed statistical analyses. *p<0.05 between groups, LSD post-hoc test.

### Reciprocal Social Interaction in Oprm1 Mutant Mice

Mendelian inheritance from Oprm1+/− parents leads to a larger number of Oprm1+/− offspring (50%), relative to Oprm1−/− (25%) or Oprm1+/+ (25%). To obtain comparable numbers of all three genotypes for assessment of social behavior, we performed parallel analysis of offspring from Oprm1+/− parents, and age-matched offspring of parents that were both Oprm1+/+ or Oprm1−/−. In the reciprocal social interaction test (described below), mice with the same genotype exhibited similar behavior regardless of breeding strategy (Figure 1D), and were therefore pooled for further analysis.

We first evaluated reciprocal social interaction between two freely moving age- and sex-matched mice: one “mutant” animal generated by the Oprm1 breeding strategies described above, and a “partner” that was either a novel mutant mouse of the same genotype or a wild-type C57Bl/6J (Figure 1E). The total time spent in social interaction was similar regardless of partner genotype (Figure S2), so data are pooled for presentation (Figure 1F). On average, the duration of interaction was lower in female mice than male mice (main effect of Sex: F_1,166_=10.54, p<0.01), but there was not a significant Genotype x Sex interaction (F_2,166_<1). Importantly, there was also a main effect of Genotype (F_2,166_=12.31, p<0.01), indicating both Oprm1+/− and Oprm1−/− mutants spent less time than Oprm1+/+ controls engaging in social interaction.

In the reciprocal social interaction test, the total interaction duration includes several qualitatively different types of social behavior [22,29]. In terms of affiliative social behaviors, there was a main effect of Genotype for nose contact (Figure 1G; F_2,129_=3.38, p=0.026) and huddling (Figure 1H; F_2,129_=6.92, p=0.001), with decreases in Oprm1−/− mutants that were more moderate in Oprm1+/− mutants, relative to Oprm1+/+ controls. In terms of investigative behaviors, there was a main effect of Genotype for following (Figure 1I; F_2,129_=8.26, p<0.01), but no significant change in the amount of other non-reciprocated social exploratory behaviors, such as anogenital sniffing or nose-flank contact (Figure 1J; F_2,129_=1.82, p=0.156). There were also no differences in social approach or memory in a three-chamber social test (Figure S3).

In addition to the reciprocal social behavior of the mutant mouse, we also quantified the social behavior of wild-type C57Bl/6J partners in each test session (Figure 1E). There was no difference in the total interaction duration as a function of the mutant partner genotype (Figure 1K; F_2,105_<1), but interesting trends emerged in the qualitative breakdown of specific types of social behavior. In terms of affiliative social behaviors, there were similar trends towards reduced nose contact and huddling, but not in following (Figure 1L-N). However, wildtype partners engaged in more non-reciprocated social exploratory behaviors with Oprm1−/− mutant partners (Figure 1O; F_2,83_=3.58, p=0.032). This result implies that wildtype mice adjust their own social behavior in response to their Oprm1 mutant partner – a novel finding that we subsequently investigated with other assays of social behavior.

### Attenuated Social Reward in Wild-Type Mice Housed with Oprm1 Mutants

Previous reports indicate Oprm1−/− mutants fail to develop a social conditioned place preference (CPP) when housed together with mice of the same genotype [21]. To extend these findings, we evaluated social CPP in Oprm1+/+, Oprm1+/− and Oprm1−/− mice (Figure 2A). Oprm1+/+ controls developed a preference for the bedding associated with group housing, versus a bedding associated with isolation housing (Figure 2B). This social CPP did not develop in Oprm1+/− and Oprm1−/− mutants, as the preference score in mutant mice was significantly reduced compared to Oprm1+/+ controls (Figure 2C; F_2,82_=7.443, p=0.001).

**Figure 2.**
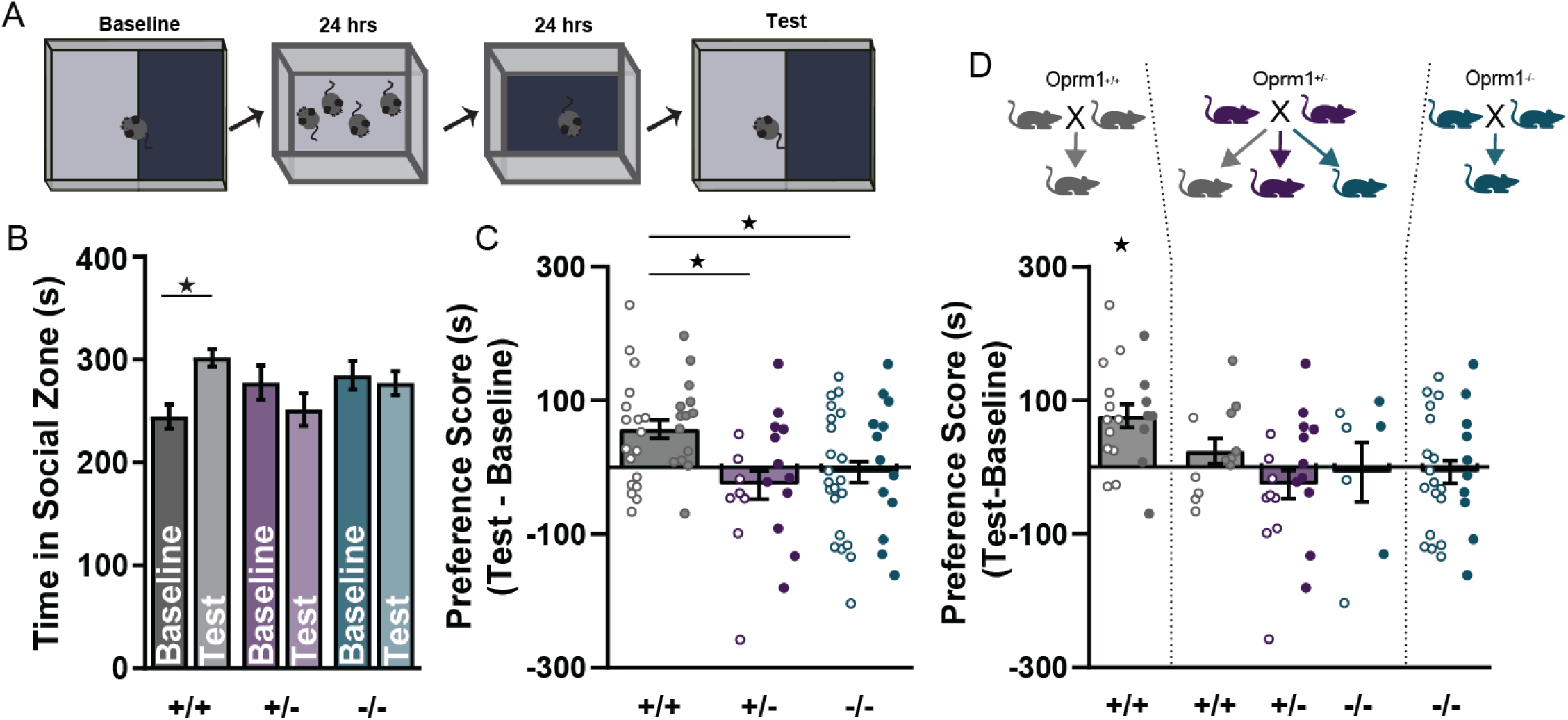
Social conditioned place preference (CPP) as a function of Oprm1 genotype and breeding strategy. **(A)** Schematic diagram of the social CPP test protocol. **(B)** Time spent in the social zone for each genotype, during the baseline session before conditioning and the test session after conditioning. **(C)** Preference scores for each genotype, calculated as time in social zone on test minus baseline. **(D)** Preference scores for each genotype, disaggregated by breeding strategy. All groups contained similar numbers of female mice (open symbols) and male mice (closed symbols); see Supplementary Table 2 for detailed statistical analyses. *p<0.05 according to paired t-test **(B)**, LSD post-hoc test **(C)**, or one-sample t-test **(D)**.

When these data were divided based on breeding strategy (Figure 2D), Oprm1+/+ mice from Oprm1+/+ parents developed a robust preference for the social bedding (F_4,76_=4.78, p=0.002). However, Oprm1+/+ mice from Oprm1+/− parents (which were conditioned with Oprm1+/− and Oprm1−/− mutant littermates) failed to develop a preference for the social bedding. This result was particularly striking because the reciprocal social interaction of Oprm1+/+ mice did not depend on parental genotype (Figure 1D), and prior literature suggests parental Oprm1 genotype does not directly influence development of social behavior in offspring [22]. Together, these findings suggest the absence of social CPP in Oprm1+/+ mice from Oprm1+/− parents is due to reduced quality of social interaction with mutant littermates during conditioning, leading to a less rewarding experience.

### Diminished Real Time Social Preference for Oprm1−/− Mutant Mice

To compare the quality of social interaction with an Oprm1 mutant mouse relative to a typical control mouse, we measured preference for typical social interaction in real time [31]. C57Bl/6J mice served as “judges” in a chamber with two confined stimulus mice (Figure 3A). One of these stimulus mice was “typical” (Oprm1+/+ wildtype), while the other stimulus mouse was “atypical” (Oprm1+/− or Oprm1−/− mutant). Both stimulus mice were age- and sex-matched to the judge. Wildtype judges failed to exhibit reliable discrimination between atypical Oprm1+/− mutants and typical Oprm1+/+ controls (Figure 3B-C). However, wildtype judges did reliably discriminate between atypical Oprm1−/− mutants and typical Oprm1+/+ controls (Figure 3D-E), exhibiting a robust social preference for the chamber containing the typical mouse (F_1,22_=5.87, p=0.002). Together, these data provide converging evidence that the abnormal social behavior exhibited by Oprm1 mutant mice can negatively influence the reciprocal social experience of conspecifics.

**Figure 3.**
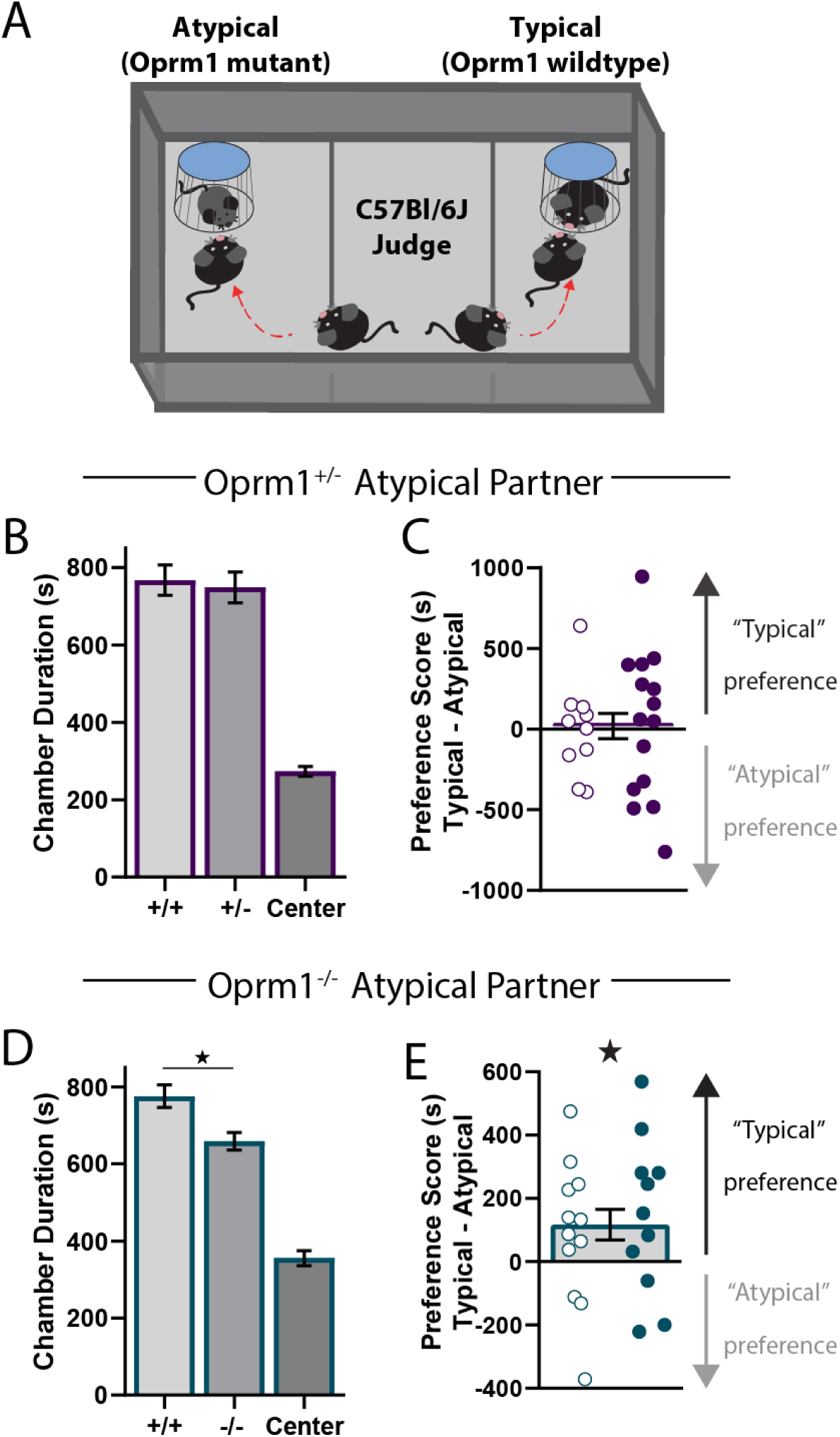
Real time social preference of genotypical judges for interaction with Oprm1 mutants. **(A)** Schematic diagram of the test procedure: genotypical judges (C57Bl/6J) simultaneously engage with social targets that are typical (Oprm1+/+) or atypical (Oprm1 mutant). **(B-C)** Time spent in each chamber **(B)** and preference score **(D)** for genotypical judges choosing between interaction with Oprm1+/+ and Oprm1+/− targets. **(D-E)** Time spent in each chamber **(D)** and preference score **(E)** for genotypical judges choosing between interaction with Oprm1+/+ and Oprm1−/− targets. All groups contained similar numbers of female mice (open symbols) and male mice (closed symbols); see Supplementary Table 2 for detailed statistical analyses. *p<0.05 according to LSD post-hoc test **(D)** or one-sample t-test **(E)**.

### Altered Striatal Microcircuitry in Oprm1 Mutant Mice

Mu opioid receptor activation in the nucleus accumbens is particularly important for social behavior [9-11], so we next assessed functional changes in synaptic transmission within the nucleus accumbens of Oprm1+/− and Oprm1−/− mutant mice. Our analysis focused on inhibitory synaptic transmission, because Oprm1−/− mutant mice have a substantial increase in symmetrical synapses within the nucleus accumbens, but no change in asymmetrical synapses [22]. To selectively analyze changes in D1- and D2-MSNs, we crossed Oprm1 knockout mice with double-transgenic fluorescent reporter mice expressing Drd1-tdTomato and Drd2- eGFP. In acute brain slices prepared from these animals, we performed whole-cell voltage-clamp recordings from red D1-MSNs and green D2-MSNs (Figure 4A-B), and measured the frequency and amplitude of miniature inhibitory postsynaptic currents (mIPSCs). In Oprm1+/+ control mice, there was a noteworthy sex difference in basal synaptic transmission, with larger mIPSC amplitude in male D1-MSNs and female D2- MSNs (Figure S4).

**Figure 4.**
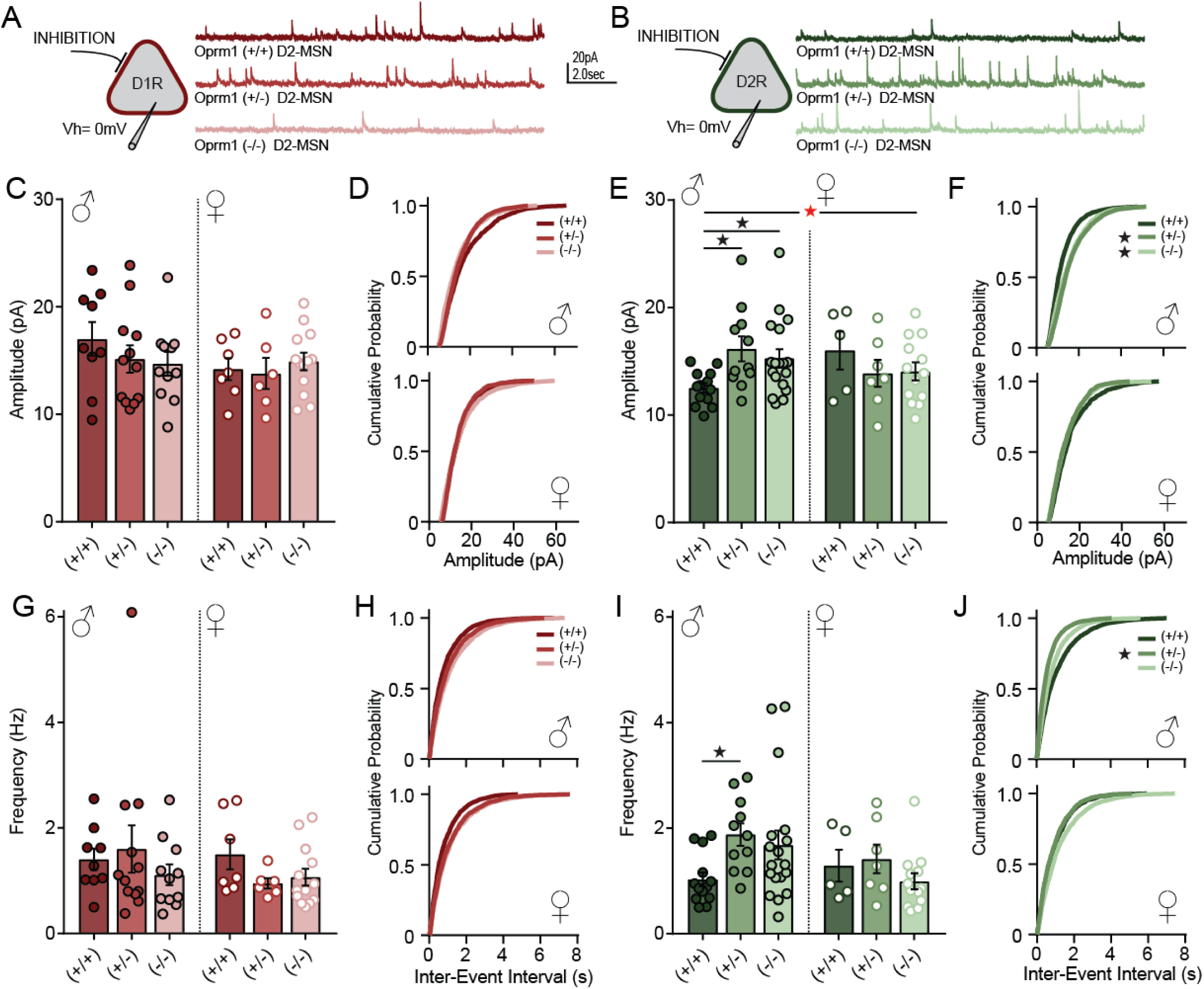
Electrophysiological recordings from medium spiny projection neurons (MSNs) in the nucleus accumbens, to assess inhibitory synaptic transmission. **(A-B)** Schematic diagram showing whole-cell voltage-clamp recordings from MSNs identified by expression of Drd1-tdTomato **(A)** or Drd2-eGFP **(B)**. Example traces show miniature inhibitory postsynaptic currents (mIPSCs) recorded for each genotype. **(C-F)** Average mIPSC amplitude and cumulative probability plots for D1-MSNs **(C-D)** and D2-MSNs **(E-F)**, separated by sex. **(G-J)** Average mIPSC frequency and cumulative probability plots for D1-MSNs **(G-H)** and D2-MSNs **(I-J)**, separated by sex. See Supplementary Table 2 for detailed statistical analyses. Red asterisk indicates a significant Genotype x Sex interaction **(E)**; *p<0.05 according to LSD post-hoc test **(E, I)** or Kolmogorov-Smirnov test comparing Oprm1 mutant to control **(F**,**J)**.

For mIPSC amplitude (Figure 4C-F), omnibus ANOVA revealed a significant Cell Type x Sex x Genotype interaction (F_2,113_=3.31, p=0.040). There were no significant effects on mIPSC amplitude in D1- MSNs (Figure 4C), but for D2-MSNs (Figure 4E), there was a significant Sex x Genotype interaction (F_2,61_=3.62, p=0.033). This interaction was driven by a main effect of Genotype in male mice (F_2,39_=4.52, p=0.017) but not in female mice (F_2,22_<1). In D2-MSNs from male mice, mIPSC amplitude was significantly higher in Oprm1+/− and Oprm1−/− mutants relative to Oprm1+/+ controls. For mIPSC frequency (Figure 4G-J), there were no significant main effects or interactions in an omnibus ANOVA (Table S2). However, we noted a trend toward a main effect of Genotype in D2-MSNs from male mice (F_2,39_=3.18, p=0.053), with higher mIPSC frequency in Oprm1+/− mutants relative to Oprm1+/+ controls.

Inhibitory synapses formed at different subcellular locations generate quantal currents with distinct biophysical properties [35,36]: perisomatic inhibitory synapses generate currents with larger amplitude, while inhibitory synapses in the dendritic arbor generate currents with smaller amplitude (Figure 5A). When we analyzed mIPSC frequency from male D2-MSNs as a function of amplitude, we found Oprm1+/− and Oprm1−/− mutants had a selective increase in the frequency of currents with amplitude larger than 10 pA (Figure 5B). Similar patterns were observed when we analyzed mIPSC frequency by either rise or decay kinetics (Figure S5), suggesting Oprm1 mutations have a selective effect on perisomatic inhibitory synapses. To visualize these synapses, we performed immunohistochemistry for gephyrin, an inhibitory postsynaptic scaffolding protein [37]. These experiments were conducted in D2-eGFP reporter mice [38], so green fluorescence could be used to construct a soma mask and quantify perisomatic gephyrin puncta (Figure 5C-D). The mean density of perisomatic gephyrin puncta was doubled in Oprm1+/− mutants, with a significant but less dramatic increase Oprm1−/− mutants (Figure 5E). Paradoxically, these structural synaptic changes did not appear to differ between sexes. Together, our results indicate that Oprm1 gene dosage alters both form and function of inhibitory microcircuits in the nucleus accumbens, which may contribute to deficits in reciprocal social behavior displayed by Oprm1 mutant mice.

**Figure 5.**
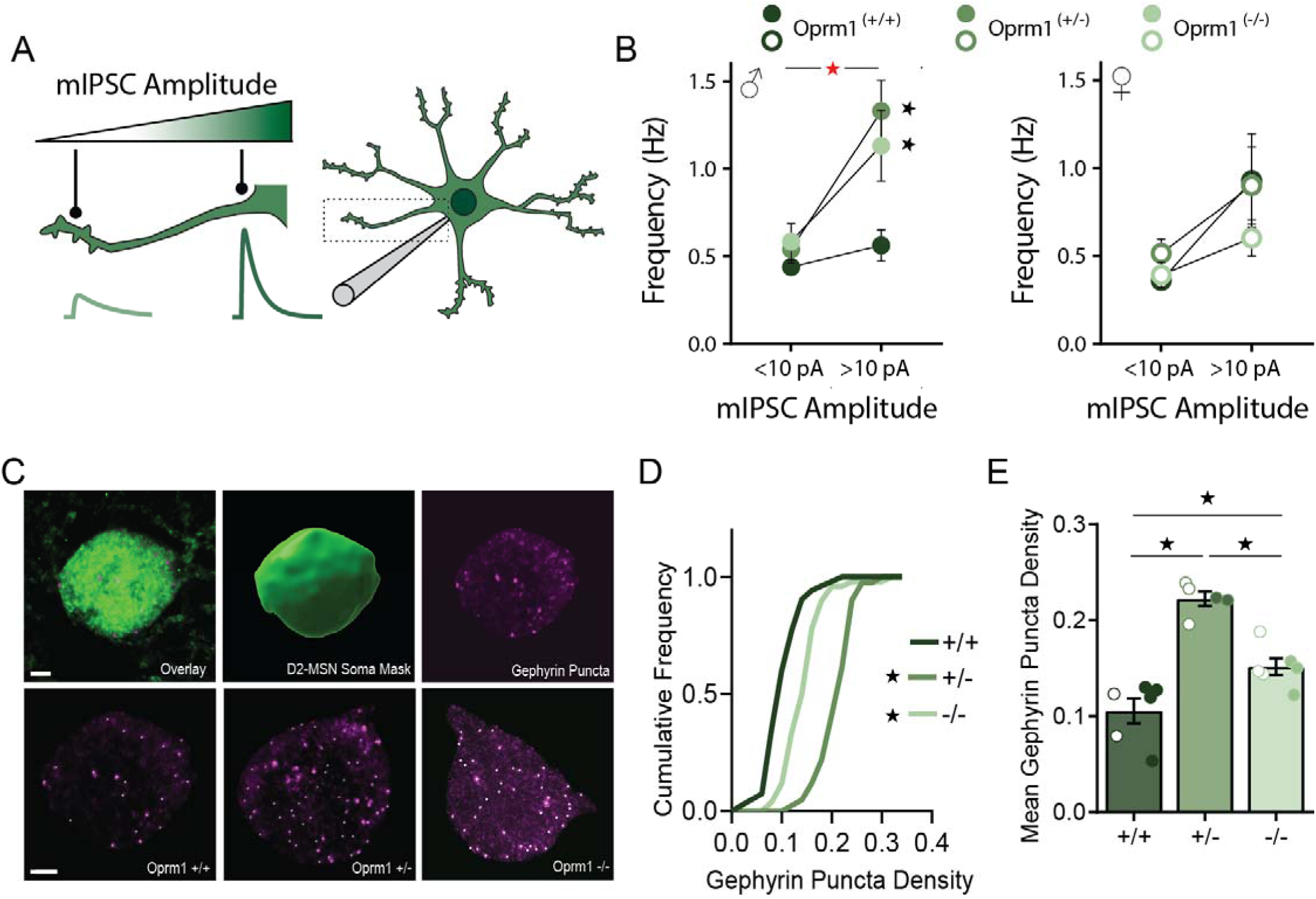
Functional and structural analysis of perisomatic inhibitory synapses in D2 medium spiny projection neurons (MSNs). **(A)** Schematic diagram showing differences in mIPSC amplitude according to location of the inhibitory synapses relative to the somatic recording electrode. **(B)** The frequency of mIPSCs with either small amplitude (<10 pA) or large amplitude (>10 pA), separated by sex. **(C)** Examples of confocal images showing D2-eGFP fluorescence (upper left) used to create a somatic mask (upper middle) for analysis of perisomatic gephyrin-immunoreactive puncta (upper right). Lower row shows representative images for each genotype, with white dots highlighting gephyrin puncta. **(D-E)** Cumulate probability plot **(D)** and mean gephyrin puncta density **(E)** for D2-MSNs from each genotype. All groups contained similar numbers of female mice (open symbols) and male mice (closed symbols); see Supplementary Table 2 for detailed statistical analyses. Red asterisk indicates a significant Genotype x Amplitude interaction **(B)**; *p<0.05 comparing Oprm1 mutant to control with LSD post-hoc test **(B)** or Kolmogorov-Smirnov test **(D)**, or LSD post-hoc test between groups **(E)**.

## DISCUSSION

Dysregulation of mu opioid receptor signaling has been reported in a variety of neuropsychiatric disorders that involve altered social behavior [15-19]. These conditions likely involve a partial (rather than complete) dysregulation of mu opioid receptor signaling, which we have modeled using mice with heterozygous genetic knockout of Oprm1. This partial reduction of mu opioid receptor signaling led to robust deficits in both reciprocal social interaction and social CPP in Oprm1+/− mice. Furthermore, the reciprocal social behavior of genotypical partners was also affected by interaction with Oprm1 mutant mice, which represents a novel aspect of social impairments caused by deficient mu opioid receptor signaling. Oprm1 gene dosage effects were evident not only in social behavior, but also at the level of inhibitory microcircuitry within the nucleus accumbens, where mu opioid receptor activation plays a particularly critical role in social behavior. Partial reductions of mu opioid receptor signaling can thus have wide-ranging impacts on both neural circuit organization and behavioral output.

### Multifaceted Influence of Oprm1 Gene Dosage on Reciprocal Social Behavior

Homozygous Oprm1 knockout mice have deficits in maternal attachment [20], social reward [21], and reciprocal social interaction [22]. We extended these analyses to Oprm1+/− mice using a breeding strategy that permitted comparison with both Oprm1+/+ and Oprm1−/− littermates. We found that Oprm1+/− mice had significant reductions in the time spent interacting with novel conspecifics in the reciprocal social interaction test, similar to the phenotype we and others observed in Oprm1−/− mice [22]. We did not detect major changes in social approach or memory in a three-chamber social test, which contrasts with a previous study of the same Oprm1 knockout mouse on a different genetic background [22]. This pattern of findings is consistent with the fact that different mouse strains vary in behavioral performance in the three-chamber social test [39], and suggest phenotypes involving reciprocal social interaction may be less sensitive to variation in background strain.

We also analyzed the behavior of genotypical partners of Oprm1 mutant mice in the reciprocal social interaction test. We found subtle indications that interaction with Oprm1 mutant mice alters the reciprocal social behavior of genotypical partners, as previously reported for other mouse strains with atypical social behavior [40]. This notion was further supported by two additional lines of evidence. First, in a test of social CPP, the preference normally observed for group housing with conspecifics was absent when genotypical mice were housed with Oprm1 mutant littermates. Second, in a test of real time social preference [31], genotypical judges exhibited a preference for interaction with other genotypical mice versus atypical Oprm1−/− mice. This preference was not observed when the atypical mouse was Oprm1+/−, so heterozygous deletion of the mu opioid receptor does not completely recapitulate all social phenotypes of homozygous Oprm1 knockout mice. Our findings are consistent with other reports that social behavior of genotypical mice can be influenced by atypical conspecifics [40-43]. This could be due to changes in the olfactory or auditory cues emitted by atypical mice [44,45], which negatively impact the quality of their interactions with atypical partners. While more research is needed to identify the specific atypical features of Oprm1 mutant mice, our data reveal new aspects of reciprocal social interaction that are influenced by deficits in mu opioid receptor signaling.

We used similar numbers of female and male animals in all tests of social behavior, and did not detect any significant interactions between sex and genotype. However, there was a main effect of sex in the reciprocal social interaction test, with female mice spending less time engaging in social interaction than male mice. This difference was most prominent in Oprm1+/− mice, although the statistical interaction between sex genotype was not significant. We note that female Oprm1+/− mice exhibited a similar trend towards lower expression of Oprm1 mRNA, pointing to the possibility of subtle sex differences arising from differential mu opioid receptor expression in male and female mice, which may be regulated by circulating sex hormones [46].

### Oprm1 Gene Dosage and Remodeling of Nucleus Accumbens Microcircuitry

To identify neurobiological correlates of altered social behavior in Oprm1 mutant mice, we focused on the nucleus accumbens. Mu opioid receptor activity in this brain region can bidirectionally modulate social behavior in rodents [9-11], and homozygous Oprm1 knockout mice show a dramatic increase in the number of symmetrical synapses within the nucleus accumbens [22]. We corroborated this prior report using gephyrin immunoreactivity as a marker of perisomatic inhibitory synapses onto D2-MSNs. The density of gephyrin puncta was significantly elevated in Oprm1−/− mice, and elevated even further in Oprm1+/− mice, with no evidence of a sex difference. This striking data show haploinsufficiency of mu opioid receptor gene expression can cause more dramatic neurobiological changes than complete genetic knockout of Oprm1, perhaps due to compensatory adaptations that occur in the total absence of mu opioid receptor expression.

In male Oprm1+/− mice, the structural reorganization of inhibitory synapses onto D2-MSNs was accompanied by altered inhibitory synaptic transmission. There was a significant increase in mIPSC amplitude and frequency in D2-MSNs from male Oprm1+/− mice, similar to previous observations in the central amygdala of male Oprm1−/− mice [47]. The increase in mIPSC frequency was particularly pronounced for events of large amplitude, which likely correspond to the perisomatic synapses detected using gephyrin immunoreactivity. Fast-spiking interneurons tend to form perisomatic inhibitory synapses with large quantal amplitude onto striatal MSNs [35], and these interneurons express the mu opioid receptor in other brain regions [48-50]. This raises the possibility that loss of mu opioid receptor expression from presynaptic neurons may contribute to remodeling of inhibitory synapses onto MSNs in male mice, although the mu opioid receptor is also expressed by postsynaptic MSNs [51,52]. Additional research is needed to determine whether inhibitory microcircuits and reciprocal social behavior are regulated by mu opioid receptor expression in specific nucleus accumbens cell types, as previously shown for responses to exogenous opioid exposure [52,53].

Paradoxically, functional changes in synaptic transmission were not observed in female Oprm1+/− mice, even though both sexes showed a comparable increase in D2-MSN gephyrin puncta density. One potential explanation for this pattern of results is that the basal mIPSC amplitude is higher in D2-MSNs of female mice and D1-MSNs of male mice. A ceiling effect may therefore have obscured our ability to detect increased mIPSC amplitude in D2-MSNs from female Oprm1 mutant mice. While sex differences at nucleus accumbens inhibitory synapses have not previously been investigated in a cell type-specific fashion, there are well-documented sex differences in the structure and function of excitatory synapses in the nucleus accumbens [54,55], including cell type-specific changes [56]. We did not evaluate excitatory synaptic transmission in this study, but the expression of genes related to glutamatergic signaling are altered in the nucleus accumbens of Oprm1−/− mice [22]. Since inhibitory synaptic transmission appeared relatively normal in female Oprm1−/− mice, changes in excitatory synaptic transmission could make a larger contribution to their atypical social behavior. However, both sexes showed robust changes in D2-MSN gephyrin puncta density, suggesting a common reorganization of inhibitory microcircuitry caused by complete or partial decrements in mu opioid receptor signaling.

### Translational Implications

Our findings demonstrate that partial disruption of mu opioid receptor signaling can have profound effects on both neural circuit organization and behavioral output. In some cases, the impact of haploinsufficiency was even greater than complete loss of mu opioid receptor signaling. The dysregulation of mu opioid receptor signaling reported in a variety of neuropsychiatric disorders may therefore reflect fundamental alterations in brain function, and contribute to the pathophysiology of these conditions [15-19]. Partial loss of mu opioid receptor signaling could be caused by genetic polymorphisms affecting the receptor itself, associated signaling proteins, and opioid peptide ligands as well as their catabolic enzymes. Conversely, genetic variants that enhance some aspects of mu opioid receptor signaling (like the Oprm1 A118G polymorphism) can increase sociability, even in the heterozygous state [57-60]. Therefore, signaling via the mu opioid receptor may not only contribute to the etiology of neuropsychiatric disorders, but also represent a target for therapeutic intervention.

## ACKNOWLEDGEMENTS

We thank Bailey Remmers and David Leipold for technical assistance, as well as Adrine Kocharian, Marc Pisansky, Cassie Retzlaff and Brian Trieu for stimulating discussions. We thank the University of Minnesota Mouse Behavior Core for use of their facilities to conduct behavioral tests, and Drs. Robert Meisel and Paul Mermelstein for generously sharing resources.

## REFERENCES

1 Panksepp J, Herman BH, Vilberg T, Bishop P, DeEskinazi FG. Endogenous opioids and social behavior. Neuroscience and Biobehavioral Reviews. 1980;4(4):473–87.

2 Trezza V, Baarendse PJJ, Vanderschuren LJMJ. The pleasures of play: pharmacological insights into social reward mechanisms. Trends in pharmacological sciences. 2010;31(10):463–9.

3 Darcq E, Kieffer BL. Opioid receptors: drivers to addiction? Nat Rev Neurosci. 2018.

4 Chelnokova O, Laeng B, Løseth G, Eikemo M, Willoch F, Leknes S. The µ-opioid system promotes visual attention to faces and eyes. Social cognitive and affective neuroscience. 2016;11(12):1902–09.

5 Achterberg EJM, van Swieten MMH, Houwing DJ, Trezza V, Vanderschuren L. Opioid modulation of social play reward in juvenile rats. Neuropharmacology. 2019;159:107332.

6 Guard HJ, Newman JD, Roberts RL. Morphine administration selectively facilitates social play in common marmosets. Dev Psychobiol. 2002;41(1):37–49.

7 Hsu DT, Sanford BJ, Meyers KK, Love TM, Hazlett KE, Wang H, et al. Response of the μ-opioid system to social rejection and acceptance. Molecular psychiatry. 2013;18(11):1211–7.

8 Hsu DT, Sanford BJ, Meyers KK, Love TM, Hazlett KE, Walker SJ, et al. It still hurts: altered endogenous opioid activity in the brain during social rejection and acceptance in major depressive disorder. Molecular psychiatry. 2015;20(2):193–200.

9 Trezza V, Damsteegt R, Achterberg EJ, Vanderschuren LJ. Nucleus accumbens mu-opioid receptors mediate social reward. J Neurosci. 2011;31(17):6362–70.

10 Resendez SL, Dome M, Gormley G, Franco D, Nevárez N, Hamid AA, et al. μ-Opioid receptors within subregions of the striatum mediate pair bond formation through parallel yet distinct reward mechanisms. Journal of Neuroscience. 2013;33(21):9140–49.

11 Smith CJW, Wilkins KB, Li S, Tulimieri MT, Veenema AH. Nucleus accumbens mu opioid receptors regulate context-specific social preferences in the juvenile rat. Psychoneuroendocrinology. 2018;89:59–68.

12 Baldo BA, Kelley AE. Discrete neurochemical coding of distinguishable motivational processes: insights from nucleus accumbens control of feeding. Psychopharmacology (Berl). 2007;191(3):439–59.

13 Richard JM, Castro DC, Difeliceantonio AG, Robinson MJ, Berridge KC. Mapping brain circuits of reward and motivation: in the footsteps of Ann Kelley. Neurosci Biobehav Rev. 2013;37(9 Pt A):1919–31.

14 Castro DC, Bruchas MR. A Motivational and Neuropeptidergic Hub: Anatomical and Functional Diversity within the Nucleus Accumbens Shell. Neuron. 2019;102(3):529–52.

15 Kennedy SE, Koeppe RA, Young EA, Zubieta JK. Dysregulation of endogenous opioid emotion regulation circuitry in major depression in women. Arch Gen Psychiatry. 2006;63(11):1199–208.

16 Prossin AR, Love TM, Koeppe RA, Zubieta JK, Silk KR. Dysregulation of regional endogenous opioid function in borderline personality disorder. Am J Psychiatry. 2010;167(8):925–33.

17 Pellissier LP, Gandía J, Laboute T, Becker JAJ, Le Merrer J. μ opioid receptor, social behaviour and autism spectrum disorder: reward matters. British Journal of Pharmacology. 2018;175(14):2750–69.

18 Ashok AH, Myers J, Reis Marques T, Rabiner EA, Howes OD. Reduced mu opioid receptor availability in schizophrenia revealed with [(11)C]-carfentanil positron emission tomographic Imaging. Nat Commun. 2019;10(1):4493.

19 Nummenmaa L, Karjalainen T, Isojarvi J, Kantonen T, Tuisku J, Kaasinen V, et al. Lowered endogenous mu-opioid receptor availability in subclinical depression and anxiety. Neuropsychopharmacology. 2020.

20 Moles A, Kieffer BL, D’Amato FR. Deficit in attachment behavior in mice lacking the mu-opioid receptor gene. Science. 2004;304(5679):1983–6.

21 Cinque C, Pondiki S, Oddi D, Di Certo MG, Marinelli S, Troisi A, et al. Modeling socially anhedonic syndromes: genetic and pharmacological manipulation of opioid neurotransmission in mice. Transl Psychiatry. 2012;2(8):e155.

22 Becker JA, Clesse D, Spiegelhalter C, Schwab Y, Le Merrer J, Kieffer BL. Autistic-like syndrome in mu opioid receptor null mice is relieved by facilitated mGluR4 activity. Neuropsychopharmacology. 2014;39(9):2049–60.

23 Matthes HWD, Maldonado R, Simonin F, Valverde O, Slowe S, Kitchen I, et al. Loss of morphine-induced analgesia, reward effect and withdrawal symptoms in mice lacking the μ-opioid-receptor gene. Nature. 1996;383(6603):822–23.

24 Shuen JA, Chen M, Gloss B, Calakos N. Drd1a-tdTomato BAC transgenic mice for simultaneous visualization of medium spiny neurons in the direct and indirect pathways of the basal ganglia. J Neurosci. 2008;28(11):2681–5.

25 Gong S, Zheng C, Doughty ML, Losos K, Didkovsky N, Schambra UB, et al. A gene expression atlas of the central nervous system based on bacterial artificial chromosomes. Nature. 2003;425(6961):917–25.

26 Lefevre EM, Pisansky MT, Toddes C, Baruffaldi F, Pravetoni M, Tian L, et al. Interruption of continuous opioid exposure exacerbates drug-evoked adaptations in the mesolimbic dopamine system. Neuropsychopharmacology. 2020.

27 Panksepp JB, Lahvis GP. Social reward among juvenile mice. Genes Brain Behav. 2007;6(7):661–71.

28 Dolen G, Darvishzadeh A, Huang KW, Malenka RC. Social reward requires coordinated activity of nucleus accumbens oxytocin and serotonin. Nature. 2013;501(7466):179–84.

29 Terranova ML, Laviola G. Scoring of social interactions and play in mice during adolescence. Curr Protoc Toxicol. 2005;Chapter 13:Unit13 10.

30 Nadler JJ, Moy SS, Dold G, Trang D, Simmons N, Perez A, et al. Automated apparatus for quantitation of social approach behaviors in mice. Genes Brain Behav. 2004;3(5):303–14.

31 Shah CR, Forsberg CG, Kang JQ, Veenstra-Vanderweele J. Letting a Typical Mouse Judge Whether Mouse Social Interactions Are Atypical. Autism Research. 2013;6(3):212–20.

32 Pisansky MT, Lefevre EM, Retzlaff CL, Trieu BH, Leipold DW, Rothwell PE. Nucleus Accumbens Fast-Spiking Interneurons Constrain Impulsive Action. Biol Psychiatry. 2019;86(11):836–47.

33 Sora I, Elmer G, Funada M, Pieper J, Li XF, Hall FS, et al. Mu opiate receptor gene dose effects on different morphine actions: evidence for differential in vivo mu receptor reserve. Neuropsychopharmacology. 2001;25(1):41–54.

34 Kieffer BL, Gaveriaux-Ruff C. Exploring the opioid system by gene knockout. Prog Neurobiol. 2002;66(5):285–306.

35 Straub C, Saulnier JL, Begue A, Feng DD, Huang KW, Sabatini BL. Principles of Synaptic Organization of GABAergic Interneurons in the Striatum. Neuron. 2016;92(1):84–92.

36 Koos T, Tepper JM, Wilson CJ. Comparison of IPSCs evoked by spiny and fast-spiking neurons in the neostriatum. J Neurosci. 2004;24(36):7916–22.

37 Tyagarajan SK, Fritschy JM. Gephyrin: a master regulator of neuronal function? Nat Rev Neurosci. 2014;15(3):141–56.

38 Gittis AH, Hang GB, LaDow ES, Shoenfeld LR, Atallah BV, Finkbeiner S, et al. Rapid target-specific remodeling of fast-spiking inhibitory circuits after loss of dopamine. Neuron. 2011;71(5):858–68.

39 Moy SS, Nadler JJ, Perez A, Barbaro RP, Johns JM, Magnuson TR, et al. Sociability and preference for social novelty in five inbred strains: an approach to assess autistic-like behavior in mice. Genes Brain Behav. 2004;3(5):287–302.

40 Yang M, Abrams DN, Zhang JY, Weber MD, Katz AM, Clarke AM, et al. Low sociability in BTBR T+tf/J mice is independent of partner strain. Physiol Behav. 2012;107(5):649–62.

41 Langford DJ, Tuttle AH, Brown K, Deschenes S, Fischer DB, Mutso A, et al. Social approach to pain in laboratory mice. Soc Neurosci. 2010;5(2):163–70.

42 Heinla I, Ahlgren J, Vasar E, Voikar V. Behavioural characterization of C57BL/6N and BALB/c female mice in social home cage - Effect of mixed housing in complex environment. Physiol Behav. 2018;188:32–41.

43 Rogers-Carter MM, Varela JA, Gribbons KB, Pierce AF, McGoey MT, Ritchey M, et al. Insular cortex mediates approach and avoidance responses to social affective stimuli. Nat Neurosci. 2018;21(3):404–14.

44 Wohr M, Roullet FI, Crawley JN. Reduced scent marking and ultrasonic vocalizations in the BTBR T+tf/J mouse model of autism. Genes Brain Behav. 2011;10(1):35–43.

45 Scattoni ML, Ricceri L, Crawley JN. Unusual repertoire of vocalizations in adult BTBR T+tf/J mice during three types of social encounters. Genes Brain Behav. 2011;10(1):44–56.

46 Loyd DR, Wang X, Murphy AZ. Sex differences in micro-opioid receptor expression in the rat midbrain periaqueductal gray are essential for eliciting sex differences in morphine analgesia. J Neurosci. 2008;28(52):14007–17.

47 Kang-Park MH, Kieffer BL, Roberts AJ, Roberto M, Madamba SG, Siggins GR, et al. Mu-opioid receptors selectively regulate basal inhibitory transmission in the central amygdala: lack of ethanol interactions. J Pharmacol Exp Ther. 2009;328(1):284–93.

48 Drake CT, Milner TA. Mu opioid receptors are extensively co-localized with parvalbumin, but not somatostatin, in the dentate gyrus. Neurosci Lett. 2006;403(1-2):176–80.

49 Glickfeld LL, Atallah BV, Scanziani M. Complementary modulation of somatic inhibition by opioids and cannabinoids. J Neurosci. 2008;28(8):1824–32.

50 Krook-Magnuson E, Luu L, Lee SH, Varga C, Soltesz I. Ivy and neurogliaform interneurons are a major target of mu-opioid receptor modulation. J Neurosci. 2011;31(42):14861–70.

51 Banghart MR, Neufeld SQ, Wong NC, Sabatini BL. Enkephalin Disinhibits Mu Opioid Receptor-Rich Striatal Patches via Delta Opioid Receptors. Neuron. 2015;88(6):1227–39.

52 Charbogne P, Gardon O, Martin-Garcia E, Keyworth HL, Matsui A, Mechling AE, et al. Mu Opioid Receptors in Gamma-Aminobutyric Acidergic Forebrain Neurons Moderate Motivation for Heroin and Palatable Food. Biol Psychiatry. 2017;81(9):778–88.

53 Cui Y, Ostlund SB, James AS, Park CS, Ge W, Roberts KW, et al. Targeted expression of mu-opioid receptors in a subset of striatal direct-pathway neurons restores opiate reward. Nat Neurosci. 2014;17(2):254–61.

54 Forlano PM, Woolley CS. Quantitative analysis of pre- and postsynaptic sex differences in the nucleus accumbens. J Comp Neurol. 2010;518(8):1330–48.

55 Meitzen J, Meisel RL, Mermelstein PG. Sex Differences and the Effects of Estradiol on Striatal Function. Curr Opin Behav Sci. 2018;23:42–48.

56 Cao J, Dorris DM, Meitzen J. Electrophysiological properties of medium spiny neurons in the nucleus accumbens core of prepubertal male and female Drd1a-tdTomato line 6 BAC transgenic mice. J Neurophysiol. 2018;120(4):1712–27.

57 Barr CS, Schwandt ML, Lindell SG, Higley JD, Maestripieri D, Goldman D, et al. Variation at the mu-opioid receptor gene (OPRM1) influences attachment behavior in infant primates. Proc Natl Acad Sci U S A. 2008;105(13):5277–81.

58 Copeland WE, Sun H, Costello EJ, Angold A, Heilig MA, Barr CS. Child mu-opioid receptor gene variant influences parent-child relations. Neuropsychopharmacology. 2011;36(6):1165–70.

59 Briand LA, Hilario M, Dow HC, Brodkin ES, Blendy JA, Berton O. Mouse model of OPRM1 (A118G) polymorphism increases sociability and dominance and confers resilience to social defeat. J Neurosci. 2015;35(8):3582–90.

60 Troisi A, Frazzetto G, Carola V, Di Lorenzo G, Coviello M, D’Amato FR, et al. Social hedonic capacity is associated with the A118G polymorphism of the mu-opioid receptor gene (OPRM1) in adult healthy volunteers and psychiatric patients. Soc Neurosci. 2011;6(1):88–97.

