## Supplemental Information for "Mu Opioid Receptor Gene Dosage Influences Reciprocal Social Behaviors and Nucleus Accumbens Microcircuitry"

### SUPPLEMENTARY MATERIALS & METHODS

#### Subjects

Female and male C57Bl/6J mice (stock #000664) and mu opioid receptor (*Oprm1*) knockout mice (stock #007559) were obtained from The Jackson Laboratory or bred in-house. As described in the text, Oprm1 mutant mice were generated using two different breeding schemes: one involving parents that were both Oprm1+/- heterozygotes (generating offspring with a mix of all possible genotypes), and another involving parents that were both either Oprm1+/+ or Oprm1-/- homozygotes (generating offspring of the same genotype). For electrophysiology and immunohistochemistry experiments, Oprm1 mutant mice were crossed with Drd1a-tdTomato reporter mice [1] and Drd2-eGFP reporter mice [2]. Mice were housed in groups of 2-5 per cage, on a 12 hour light cycle (0600h – 1800h) at ~23° C with food and water provided ad libitum. Experimental procedures were conducted between 1000h – 1600h.

#### Quantitative RT-PCR

For quantitative RT-PCR, tissue was snap frozen on dry ice and stored at -80°C. RNA was isolated using the RNeasy Mini Kit (Qiagen) according to the manufacturer's instructions. All RNA samples had A260/A280 purity ratio ≥ 2. Reverse transcription was performed using Superscript III (Invitrogen). For each sample, duplicate cDNA preparations were set up. Mouse β-actin mRNA was used as the endogenous control. Quantitative RT-PCR using SYBR green (BioRad, Hercules, CA) was carried out with a Lightcycler 480 II (Roche) system with the following cycle parameters: 1 x (30 sec @ 95°C), 35 x (5 sec @ 95°C followed by 30 sec @ 60°C). Data were analyzed by comparing the C(t) values of the treatments tested using the ΔΔC(t) method. Expression values of target genes were first normalized to the expression value of β-actin. The median of each cDNA replicate reactions was used to quantify the relative target gene expression. Primers were designed in Primer3 and validated in BLAST and are listed in Table S1.

**Behavioral Responses to Morphine Administration**

We tested open-field locomotor activity in a clear plexiglass arena (ENV-510, Med Associates) housed within a sound-attenuating chamber. The location of the mouse within the arena was tracked in two dimensions by arrays of infrared beams, connected to a computer running Activity Monitor software (Med Associates). Mice were habituated to the chamber for one hour the day before initiating drug treatment, the next day animals were administered saline and tested in the open field chamber, they were then tested on the following doses of morphine (2.0, 6.32, 20 mg/kg), receiving an incremental increase in dose every day.

Thermal antinociception was tested on a 55°C hot plate (IITC Life Scientific). The day before initiating drug treatment, mice were habituated to the instrument for 60 seconds at room temperature. We then established baseline latency to either jump or lift and lick a hind paw at 55°C. Mice were then tested 30 minutes after injection of saline or morphine, with a maximal cutoff of 30 seconds to prevent tissue damage. The percent maximum possible effect was calculated as (test latency – baseline latency) / (30 sec – baseline latency) x 100.

**Assays of Social Behavior**

Animals were moved to an isolated testing room 1 hour before tests of social behavior. All experiments were conducted at 60-70 luminosity, and at temperature conditions equal to those of the animal housing facility. Experimental sessions were video recorded and, in the case of three-chamber and social CPP, behavioral data was analyzed using ANY-maze behavioral tracking software. Dyadic social interaction was hand scored by researchers blind to experimental conditions.

<u>Reciprocal social interaction test</u>: mice were tested at 6-8 weeks of age in an opaque white rectangular box with 1 cm of fresh corn cob bedding on the floor. Experimental mice (Oprm1 mutants) were introduced to age- and sex-matched stimulus mice in the testing apparatus for 10min. Stimulus mice were either C57Bl/6J wildtype mice, or Oprm1 mutants with the same genotype as the experimental mouse [3], with similar results under both conditions (Figure S2). Videos recordings were scored using the ButtonBox software to track the frequency and duration of various social behaviors of either experimental mice or C57Bl/6J stimulus mice. Social behaviors were categorized into one of the following groups: nose-nose interaction, huddling, social exploration

(any direct contact outside of the oral-facial area), and following. A small number of videos were lost before these specific behaviors could be scored, resulting in a smaller sample size compared to total interaction duration.

Three-chamber social test: mice were tested at 6-8 weeks of age in a white plastic rectangular box (25" x 15"x 8") consisting of three interconnected chambers. Two identical wire cups were placed on each end of the apparatus. Prior to testing, mice were habituated to the empty apparatus for 10 minutes of free exploration. During the sociability test, an age- and sex-matched C57Bl/6J stimulus mouse was introduced in one wire cup, whereas the other cup was left empty. The experimental mouse was then allowed to freely explore all three chambers for ten minutes. For the social memory portion of the test, a novel age- and sex-matched C57Bl/6J stimulus mouse was introduced into the previously empty wire cup. The experimental mouse was then allowed to freely explore all three chambers for ten minutes. All three ten-minute sessions were recorded by a video camera, and time spent by the experimental mouse in each chamber and in proximity of each cylinder (<2 cm) was measured by ANY-maze tracking software. After each test, the entire apparatus was cleaned with 70% ethanol.

Social conditioned place preference (CPP): mice were weaned at 3 weeks of age into 'home' cages containing 3-5 cage-mates and housed on corn-cob bedding. Mice in each cages were tested as a group for social CPP on week later. The CPP test apparatus (18" x 10" x 8") was divided into two equally sized zones by a clear plastic wall, with a (2" x 1.5") oval opening at the base. The floor of each zone was covered with a different type of novel bedding (cellunest or small animal pellet bedding, PetSmart). The protocol began with a baseline CPP test, with each mouse allowed to freely explore the test apparatus for 10 minutes. Behavior was video-recorded and time spent in each zone was analyzed automatically using ANY-maze behavioral tracking software. After establish baseline preference for the two different beddings, mice were assigned to receive social conditioning (with cage-mates) for 24 hours on one type of bedding, followed by 24 hours in isolation on the other type of bedding. The assignment of each bedding to social or isolation conditioning was counterbalanced for an unbiased design. After isolation conditioning, animals were returned to the CPP apparatus for a 10 minutes test session. A "preference score" was calculate by taking difference between time spent in the social zone on test versus baseline.

Real-time social preference test: this assay was based on a published protocol that allows a genotypical "judge" to choose between interacting with a "typical" (Oprm1+/+) and an "atypical" (Oprm1 mutant) mouse [4].

It was conducted when mice were 6-8 weeks old, using the same three-chamber social testing apparatus described above. The genotypical judges (C57B/6J wildtype) were habituated for 10 minutes prior to testing in the empty apparatus. After habituation, two wire cups were placed in either end chamber: one contained a Oprm1+/+ mouse, and the other contained either a Oprm1+/- or Oprm1-/- mutant. Judges were then allowed to freely explore the chamber for 30 minutes. Test sessions were recorded by a video camera and the time the target mouse spent in each chamber and in proximity of each cylinder (<2 cm) was measured by ANY-maze tracking software. After each test, the entire apparatus was cleaned with 70% ethanol.

**Electrophysiology**

Parasagittal slices (240 µm) containing the nucleus accumbens were prepared from Oprm1+/+, Oprm1+/-, and Oprm1-/- mice carrying the Drd1-tdTomato or Drd2-eGFP reporter gene. Mice were anesthetized with isoflurane and decapitated, brains quickly removed and placed in ice-cold cutting solution containing (in mM): 228 sucrose, 26 NaHCO3, 11 glucose, 2.5 KCl, 1 NaH2PO4-H2O, 7 MgSO4-7H20, 0.5 CaCl2-2H2O. Slices were cut by adhering the lateral surface of the brain to the stage of a vibratome (Leica VT1000S), and allowed to recover for a minimum of 60 min in a submerged holding chamber (~25°C) containing artificial cerebrospinal fluid (aCSF) containing (in mM): 119 NaCl, 26.2 NaHCO3, 2.5 KCl, 1 NaH2PO4-H2O, 11 glucose, 1.3 MgSO4-7H2O, 2.5 CaCl2-2H2O. Slices were transferred to a submerged recording chamber and continuously perfused with aCSF at a rate of 2 mL/min at room temperature. All solutions were continuously oxygenated (95% O2/5% CO2). To pharmacologically isolate mIPSCs, D-APV (50 mM) and NBQX (10 mM) were added to block NMDARs and AMPARs, respectively and TTX (0.5 µM) to block spontaneous activity.

Whole-cell recordings from MSNs in the NAc shell were obtained under visual control using IR-DIC optics on an Olympus BX51W1 microscope. Red and green fluorescence were used to identify D1-MSNs and D2-MSNs, respectively. Voltage-clamp recordings were made with borosilicate glass electrodes (2–5 MU) filled with (in mM) 120 CsMeSO4,15 CsCl, 10 TEA-Cl, 8 NaCl, 10 HEPES, 1 EGTA, 5 QX-314, 4 ATP-Mg, 0.3 GTP-Na (pH 7.2-7.3). MSNs were voltage clamped at 0 mV to increase the driving force for current flow through $GABA_A$ receptors. Recordings were performed using a MultiClamp 700B (Molecular Devices), filtered at 2 kHz, and digitized at 10 kHz. Data acquisition and analysis were performed online using Axograph software. Series resistance was monitored continuously and experiments were discarded if resistance changed by >20%. At least

200 events per cell were acquired in 15 s blocks and detected using a threshold of 5 pA; all events included in the final data analysis were verified by eye.

#### Immunohistochemistry

Oprm1+/+, Oprm1+/-, and Oprm1-/- mice carrying the Drd2-eGFP reporter gene were deeply anesthetized using sodium pentobarbitol (Fatal-Plus, Vortech Pharmaceuticals) and transcardially perfused with ice cold 0.01 M PBS followed by ice cold 4% PFA in 0.01 M PBS.. Brains were removed and post-fixed 24 hours in 4% PFA in PBS. The following day, brains were rinsed briefly with 0.01 M PBS and sectioned at 50 um. Tissue sections were blocked for 1 hour in blocking buffer (2% NHS, 0.2% triton x 100, and 0.05% Tween20 in 0.01 M PBS) and exposed to rabbit anti-GFP (1:1000, Abcam) and mouse anti-Gephyrin (1:250, Synaptic Systems) diluted in blocking buffer. After 24 hours at 4° C, sections were rinsed in wash buffer (Tris-buffered saline with 0.1% Tween20) and exposed to anti-Rabbit A488 and anti-Mouse A647 secondary antibodies (1:1000, Abcam) overnight at 4° C.

#### Confocal Microscopy

Stained tissue sections were imaged on a Leica TCS SPE laser scanning confocal microscope (Leica Microsystems, Wetzlar, Germany). A minimum of 3 image stacks per hemisphere were collected from D2-MSNs in the nucleus accumbens of each section, including both core and shell subregions. Image stacks were collected with a Leica 63X HCX PL APO objective with numerical aperture of 1.4, using laser and PMT settings optimized for excitation and emission of Alexa A488 and A647. Digital zoom between 8x and 10x was applied and stacks were collected at 2048 by 2048 pixel resolution using a step size of 0.3 µm and 1 airy unit pinhole diameter. Image stacks were imported into Imaris 9.0 (Bitplane, Zurich, Switzerland) and analyses were conducted on 3D renderings of compiled confocal stacks. A surface object was applied to the A488 channel to produce a surface representing the GFP-expressing somata in the image stack. Using this surface as a mask, the portion of the A647 channel contained within this surface was isolated to restrict our analysis to individual D2-MSNs. The spot detection algorithm [5] was used to detect gephyrin puncta in the masked A647 channel. A second algorithm was applied to restrict spots within 1 µm of the GFP immunoreactive surface object. Puncta area density was calculated as the ratio of detected A647 spots to area of the surface object.

**Statistical Analyses**

Similar numbers of male and female animals were used in all experiments. Individual data points from males (filled circles) and females (open circles) are distinguished in figures. Sex was included as a variable in factorial ANOVA models analyzed using IBM SPSS Statistics v24, with repeated measures on within-subject factors, but sex effects were not significant unless noted otherwise. For main effects or interactions involving repeated measures, the Huynh-Feldt correction was applied to control for potential violations of the sphericity assumption. This correction reduces the degrees of freedom, resulting in non-integer values. Significant interactions are indicated in figures by a red asterisk, and were decomposed by analyzing simple effects (i.e., the effect of one variable at each level of the other variable). Significant main effects were analyzed using LSD post-hoc tests. Effect sizes are expressed as partial eta-squared ($\eta_p^2$) values. The Type I error rate was set to $\alpha$=0.05 (two-tailed) for all comparisons.

### SUPPLEMENTARY FIGURES & FIGURE LEGENDS

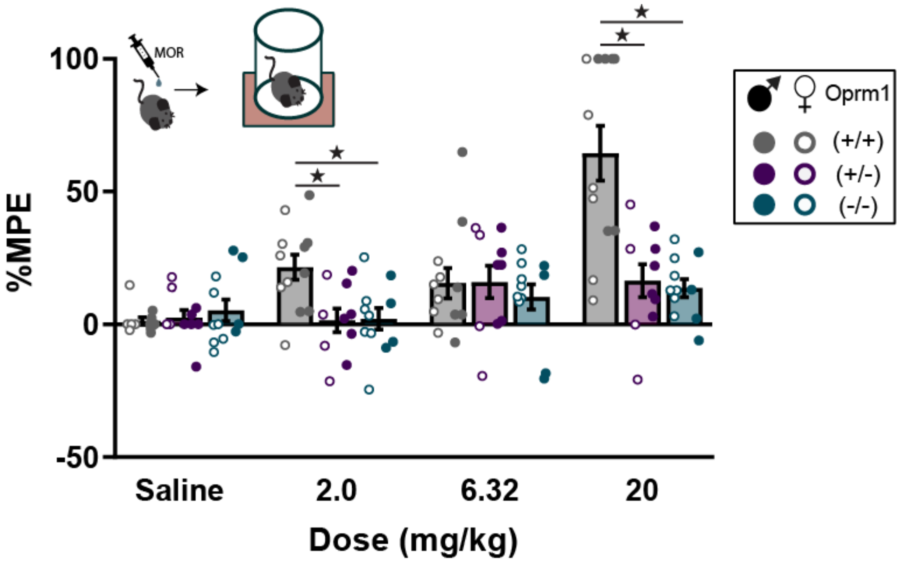


**Figure S1. Thermal antinociception after morphine administration in mu opioid receptor (Oprm1) mutant mice.** Percent maximum possible effect (% MPE) on the hot plate after injection of morphine is shown for Oprm1+/+, Oprm1+/-, and Oprm1-/- mice. All groups contained similar numbers of female mice (open symbols) and male mice (closed symbols); see Supplementary Table 2 for detailed statistical analyses. *p<0.05 between groups, LSD post-hoc test.

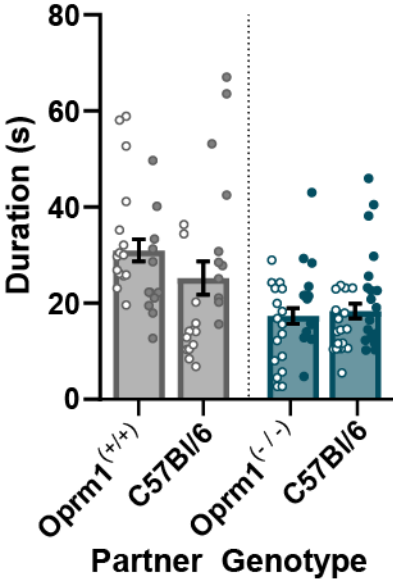


**Figure S2. Reciprocal social interaction as a function of social partner.** Total interaction time of Oprm1+/+ (grey) and Oprm1-/- (blue) mice was similar with a social partner of the same genotype, or a C57Bl/6J social partner. All groups contained similar numbers of female mice (open symbols) and male mice (closed symbols); see Supplementary Table 2 for detailed statistical analyses.

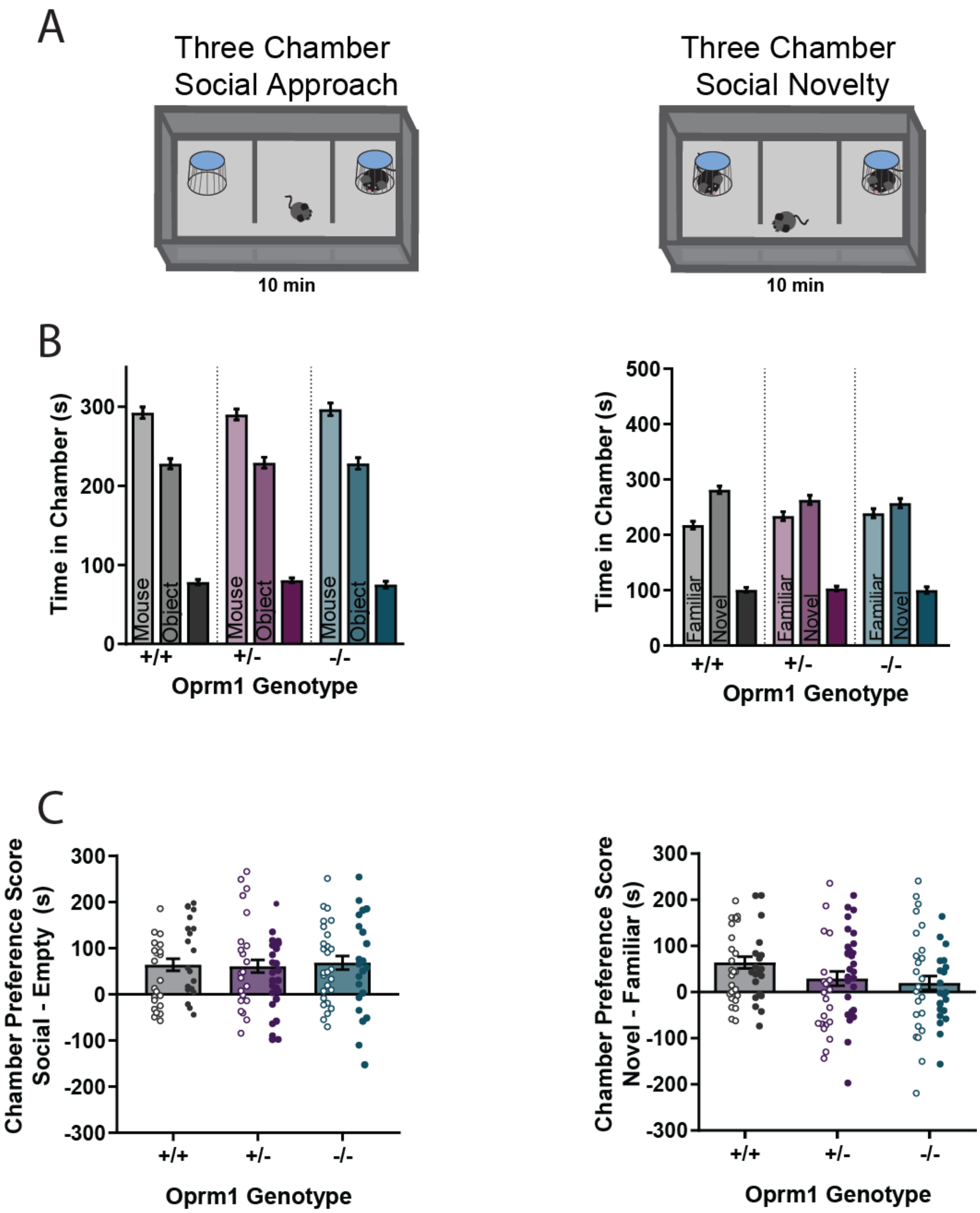


**Figure S3. Three-chamber test of social approach and memory. (A)** Schematic diagram of the three-chamber test, including conditions for evaluating social approach (*left*) and social memory (*right*). **(B)** Time spent in each chamber during social approach (*left*) and social memory (*right*). **(C)** Preference scores for social approach (*left*) and social memory (*right*). All groups contained similar numbers of female mice (open symbols) and male mice (closed symbols); see Supplementary Table 2 for detailed statistical analyses.

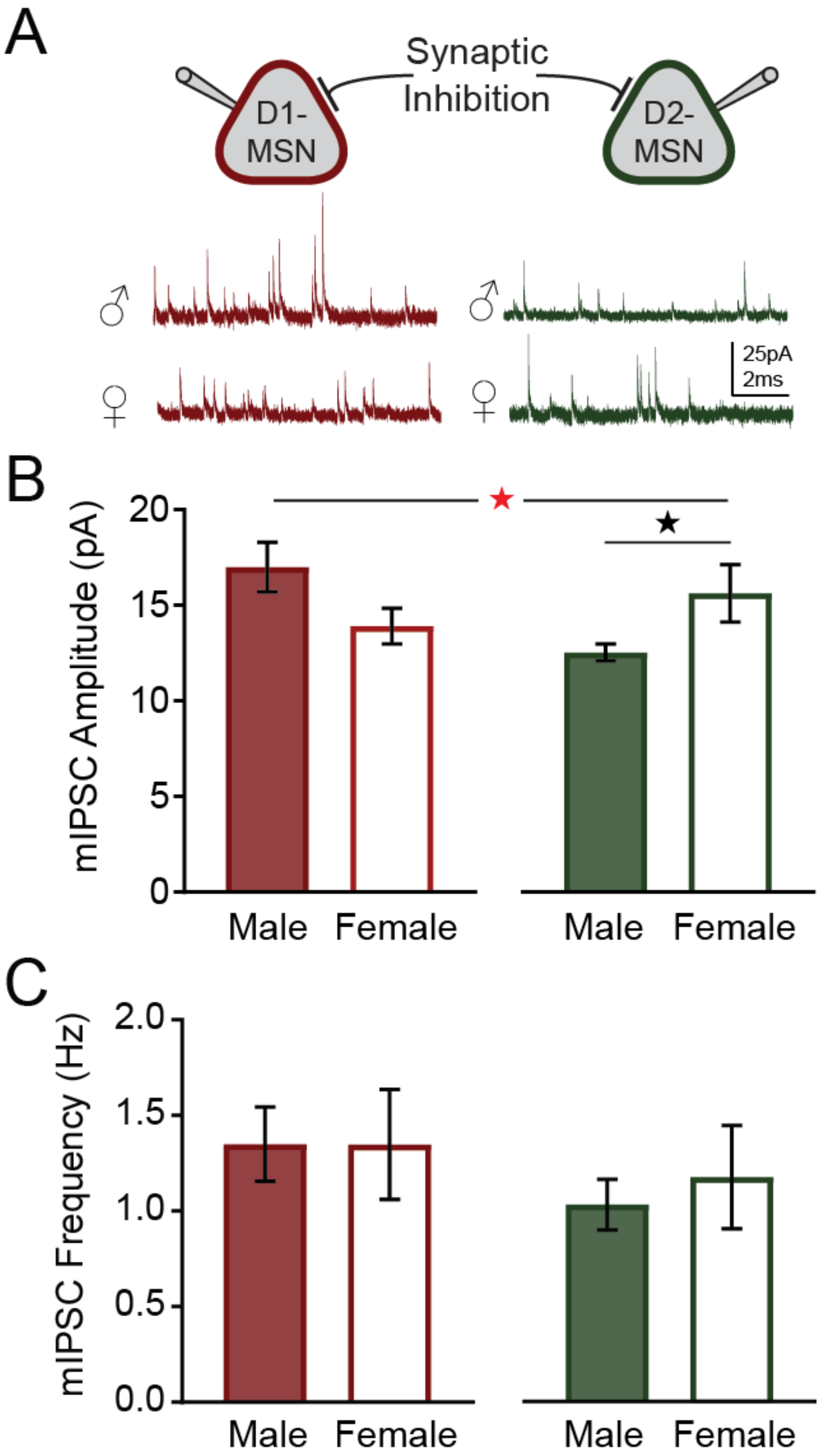


**Figure S4. Basal inhibitory synaptic transmission in male and female medium spiny neurons (MSNs). (A)** Schematic diagram of whole-cell voltage-clamp recordings from D1-MSNs (red) and D2-MSNs (green) in genotypical Oprm1+/+ mice, along with example traces from cells in male and female animals. **(B)** Average mIPSC amplitude from each cell type in male and female animals. **(C)** Average mIPSC frequency from each cell type in male and female animals. Red asterisk indicates a significant Cell Type x Sex interaction **(B)**; see Supplementary Table 2 for detailed statistical analyses. *p<0.05 between groups, simple effect test.

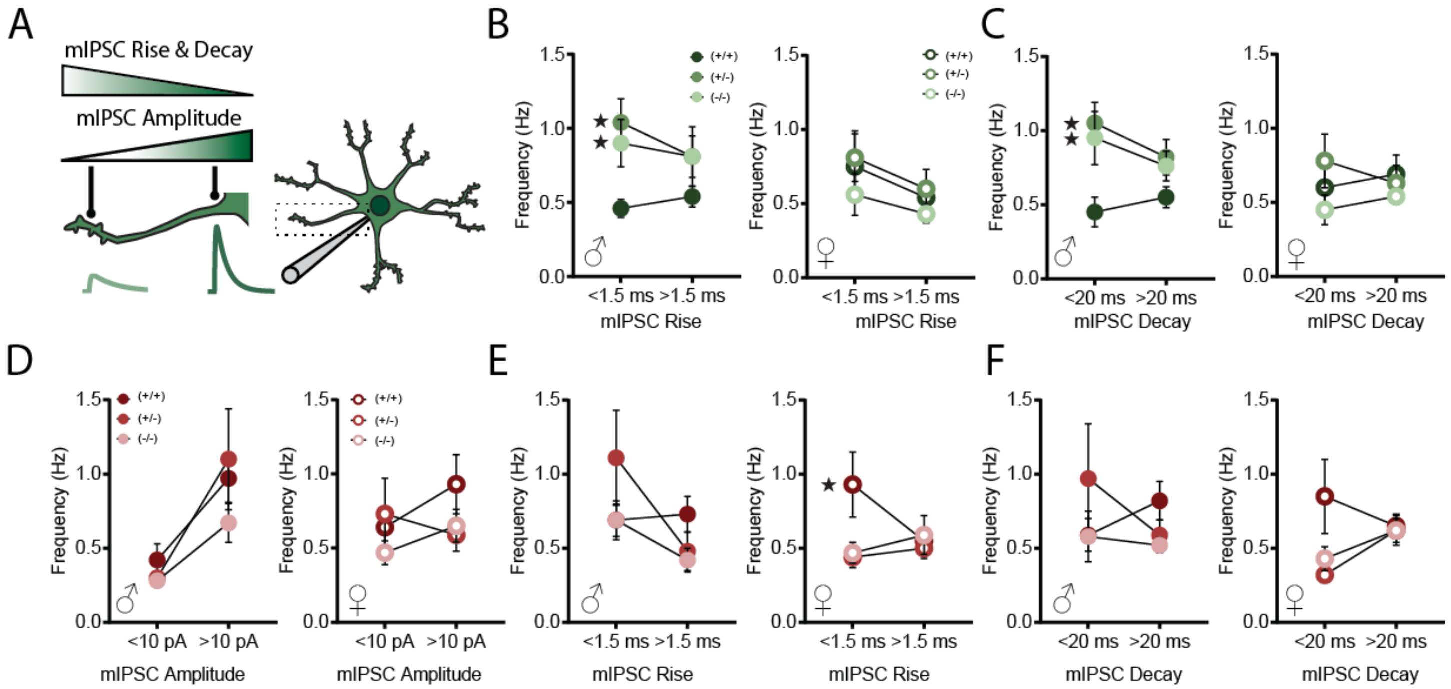


**Figure S5. Characteristics of mIPSCs recorded from male and female MSNs. (A)** Schematic diagram showing differences in mIPSC amplitude, rise time, and decay kinetics according to location of the inhibitory synapses relative to the somatic recording electrode. **(B-C)** Average mIPSC frequency in D2-MSNs in each genotype and sex, as a function of rise time **(B)** and decay kinetics **(C)**. **(D-F)** Average mIPSC frequency in D1-MSNs in each genotype and sex, as a function of amplitude **(D)**, rise time **(E)** and decay kinetics **(F)**. *p<0.05 comparing Oprm1 mutants to wild-type littermates using LSD post-hoc test; see Supplementary Table 2 for detailed statistical analyses.

**Table S1:** List of primers used for quantitative RT-PCR.

| Gene Name | Symbol | Forward oligonucleotide | Reverse oligonucleotide |
|---|---|---|---|
| beta-actin | β-actin | GACGGCCAGGTCATCACTAT | CCACCGATCCACACAGAGTA |
| Mu opioid receptor | Oprm1 | GTCACAGCCATCACCATCA | GCCAGAGCAAGGTTGAAAATG |

### SUPPLEMENTARY REFERENCES


1 Shuen JA, Chen M, Gloss B, Calakos N. Drd1a-tdTomato BAC transgenic mice for simultaneous visualization of medium spiny neurons in the direct and indirect pathways of the basal ganglia. J Neurosci. 2008;28(11):2681-5.

2 Gong S, Zheng C, Doughty ML, Losos K, Didkovsky N, Schambra UB, et al. A gene expression atlas of the central nervous system based on bacterial artificial chromosomes. Nature. 2003;425(6961):917-25.

3 Becker JA, Clesse D, Spiegelhalter C, Schwab Y, Le Merrer J, Kieffer BL. Autistic-like syndrome in mu opioid receptor null mice is relieved by facilitated mGluR4 activity. Neuropsychopharmacology. 2014;39(9):2049-60.

4 Shah CR, Forsberg CG, Kang JQ, Veenstra-Vanderweele J. Letting a Typical Mouse Judge Whether Mouse Social Interactions Are Atypical. Autism Research. 2013;6(3):212-20.

5 Banovic D, Khorramshahi O, Owald D, Wichmann C, Riedt T, Fouquet W, et al. Drosophila neuroligin 1 promotes growth and postsynaptic differentiation at glutamatergic neuromuscular junctions. Neuron. 2010;66(5):724-38.
